# Effect of *Lactobacillus reuteri* on intestinal microflora and immune parameters: involvement of sex differences

**DOI:** 10.1101/312678

**Authors:** Jiayi He, Lingyi Wu, Zhen Wu, Daodong Pan, Yuxing Guo, Xiaoqun Zeng, Liwei Lian

**Author notes:** Corresponding authors: Zhen Wu, Key Laboratory of Animal Protein Food Deep Processing Technology of Zhejiang Province, Ningbo University. Co-corresponding author: Daodong Pan, Key Laboratory of Animal Protein Food Deep Processing Technology of Zhejiang Province, Ningbo University.

## Abstract

Probiotic candidate *L. reuteri* was screened out for *in vivo* experiments based on a relatively higher gastrointestinal tolerance and moderate adhesiveness. As results shown in *in-vivo* experiments, a significantly higher level of IL-12 at low-dose group was found both in females and males. Higher levels of T-lymphocytes were also observed in females compared to control group, however, males displayed a reduction expcept for CD8-positive cells in ileum. In comparison to the control group, the relative abundance of phylotypes in the phylum *Bacteroidetes* (genus of *Bacteroides*, *Prevotella*) and *Firmicutes* (genus of *ClostridiumIV*) exihibited a reserve shift between sexes after *L. reuteri* intervened. Meanwhile, the relative abundance of several taxa (*Acetobacteroides*, *Lactobcaillus*, *bacillus*) also differed markedly in sexes at low-dose group, together with microbiota diversity, as indicated by Shannon index.

**Importance:** Sexual dimorphism has triggered researchers’ attention. However, the relationship between immune parameters and gut microbiota caused by *Lactobacillus* at different dosage are not fully elucidated. In present research, the possible probiotic role of *L. reuteri* DMSZ 8533 on immunomodulation and effect on fecal microbiota composition were investigated. Our findings demonstrate the importance of L. reuteri DMSZ 8533 as a potential probiotic strain with an immunomodulatory effect, which also alters the microflora composition depending on the sex of the host.

## Introduction

*Lactobacillus*, which are generally considered probiotic, are the resident microflora in the human gastrointestinal tract (GIT). They exhibit various health-promoting effects, including anti-cancer [1,2] and anti-oxidation [3] effects, while helping to maintain microflora balance [4] and assisting in immuno-regulation [5]. Many *Lactobacillus* species are considered safe (GRAS) [6] and food-grade. They are involved in the fermentation of food such as in dry-fermented sausage [7] and yoghurt [8] among others.

To exert beneficial effects on a host, *Lactobacillus* must survive in, and colonize, the human GIT. Therefore, a high tolerance to low pH and bile toxicity are indispensable for achieving any potential probiotic effects [9,10]. Some *Lactobacillus* have survived in an adequate dose after passing through the human GIT *in vitro*. However, some strains displayed poor survival [11,5]. Adhesion ability is a key criteria, which has mainly been evaluated by performing *in vitro* experiments [12,13,14,15]. The HT-29 and Caco-2 cell lines are widely accepted models for the evaluation of bacterial adherence due to the fact that their morphological and functional properties mimic those of mature enterocytes [16].

Oral treatment with *Lactobacillus* has been shown to enhance immune response at the systemic [17,18] and mucosal [19] levels. Interaction between *Lactobacillus* and the intestinal mucosal system exerts vital effects on gut-associated immunoglobulin secretion, and on CD4 and CD8 T-lymphocyte activation [20]. Furthermore, *Lactobacilli* stimulate CD4-positive cells [21], which can differentiate into T helper type 1 or 2 (Th1/Th2) lymphocytes. Th1 cells mainly mediate cellular immunity associated with the cytokines Interferon-γ and TNF-α, while Th2 cells drive the humoral immune response by secreting IL-4 and IL-5. However, the *Lactobacillus-*influenced immunomodulatory effect is strain-specific [22,23], and also is related to the concentration of *Lactobacillus* [24].

In addition to immune effects, the ingestion of “probiotic *Lactobacillus*” may modify and balance intestinal microflora by competing with pathogenic microorganisms [25] and by maintaining favorable microflora [26]. The Illumina MiSeq platform based on 16S rRNA gene amplicons has been used to investigate human gut microflora, including the composition of intestinal microflora before and after probiotic intake [27].

Recently, sexual dimorphism has triggered researchers’ interest. Studies have found sex-associated differences in relation to the immune system [28], gut community composition [29]and metabolic activity [30]. Sex differences in relation to immune parameters and gut microbiota caused by *Lactobacillus* have not yet been fully elucidated. The objective of this study is to evaluate and screen out a comparatively more tolerant and adhesive *Lactobacillus* strain among three *Lactobacillus* (*plantarum* 14917, *acidophilus* ATCC 4356 and *reuteri* DMSZ 8533) *in vivo*. Furthermore, sex differences in immunomodulatory effects and microflora composition were investigated in a BALB/c mice model.

## Results

### Tolerance and adhesion-related property of *Lactobacillus* strains

As shown in Fig. 1A, *L. reuteri* exhibited a high tolerance to low pH and bile salts, retaining its viability with only a 1.8 log reduction. In comparison, the viable numbers for *L. plantarum* and *L. acidophilus* decreased dramatically (5.28 logs and 2.98 logs, respectively). Meanwhile, all tested *Lactobacillus* strains showed high aggregation values ranging from 20.40% to 26.03% (Fig. 1B). However, no significant difference (P > 0.05) was found among tested strains. The percentage of cell surface hydrophobicity for *L. plantarum*, *L. acidophilus* and *L. reuteri* was significantly different (P >0.05), and *L. reuteri* was characterized by the highest affinity to xylene and a modest adhesion value (3.82%). However, *L. plantarum* and *L. acidophilus* both exhibited low cell surface hydrophobic values (less than 20%) and limited adhesive values (Fig. 1C and 1D).

**Fig 1.**
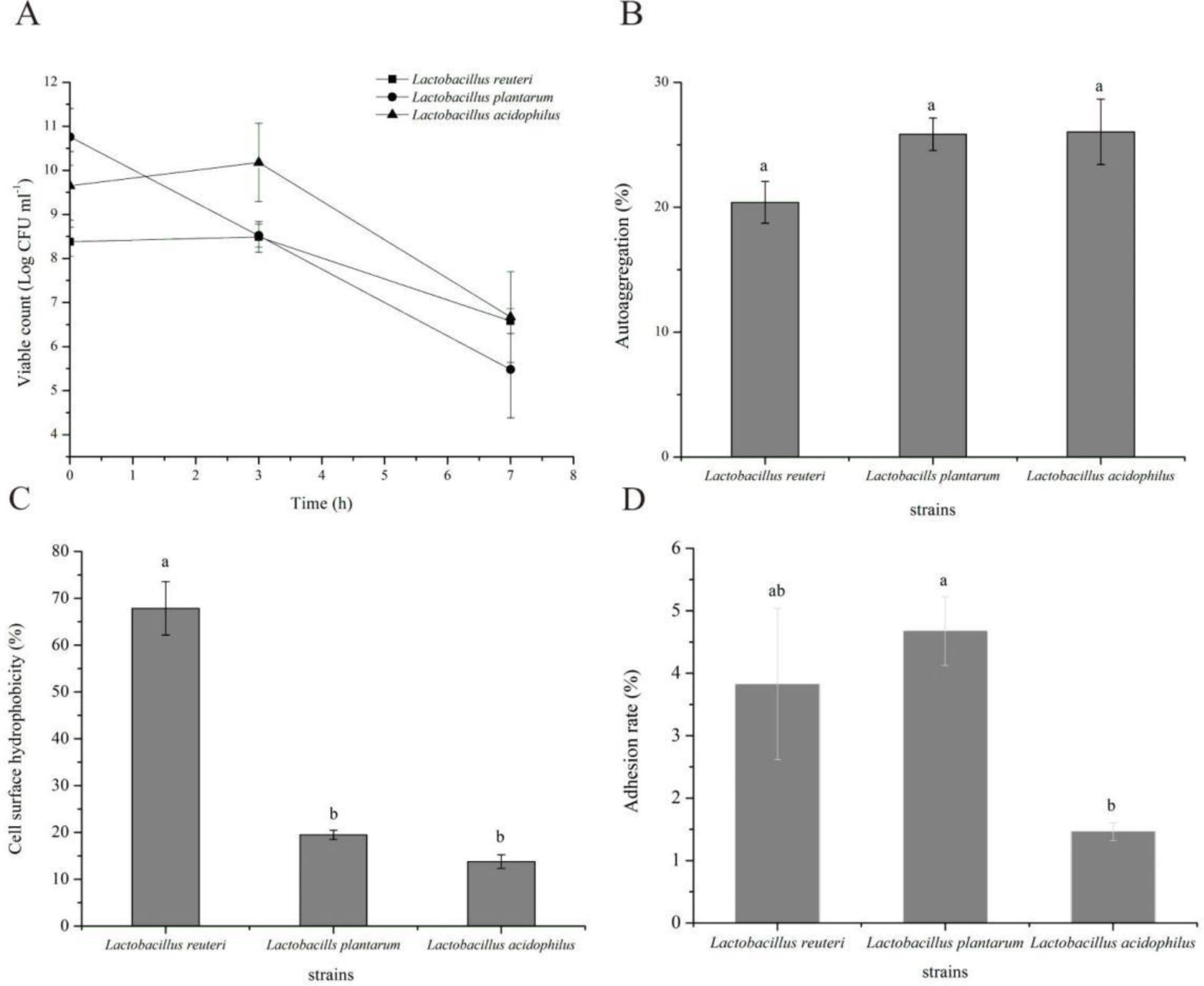
Screening experiment of probiotic *Lactobacillus*. (A) Survivability of *Lactobacillus* strains after low pH (pH=2.0) and bile salt (0.5%) treatment; (B) Auto-aggregation analysis. “a” means the index did not differ significantly (P>0.05, n=4); (C) Cell surface hydrophobicity analysis. “a” and “b” mean the indices differed significantly (P>0.05, n=4); (D) Adhesion to Caco-2 cell line. “a” and “b” mean the indices differed significantly (P >0.05, n =3). All data are presented as mean ± SE.

### Organ mass and Cytokine Measurement in BALB/c mice

As Table 1 shows, treatment with a low dose of *L. reuteri* significantly decreased the spleen index compared with males in the high-dose group, although there was a slight increase observed in females. *L. reuteri* treatment (low- and high-dose groups) significantly decreased the thymus index compared with control group males, while there was no significant difference among females in the groups. Treatment with high doses of *L. reuteri* significantly decreased the liver index compared with males and females in the control group.

**Table 1.** Effect of *L. reuteri* DMSZ 8533 on immune organ index

| group | spleen index (%) |  | thymus index (%) |  | liver index (%) |  |
| --- | --- | --- | --- | --- | --- | --- |
|  | female | male | female | male | female | male |
| control | 0.480±0.033 | 0.369±0.018<sup>ab</sup> | 0.261±0.059 | 0.166±0.024<sup>a</sup> | 4.890±0.148<sup>a</sup> | 4.954±0.334<sup>a</sup> |
| low dose | 0.482±0.013 | 0.336±0.021<sup>b</sup> | 0.267±0.031 | 0.120±0.012<sup>b</sup> | 4.874±0.158<sup>a</sup> | 4.669±0.241<sup>ab</sup> |
| high dose | 0.487±0.024 | 0.371±0.032<sup>a</sup> | 0.219±0.057 | 0.134±0.026<sup>b</sup> | 4.646±0.177<sup>b</sup> | 4.472±0.170<sup>b</sup> |

A) Values in a column with different superscript letters mean the index differ significantly ( $P < 0.05$ , $n = 6$ ).

The IL-12 and TNF-α cytokine production profiles induced by different doses of *L. reuteri* are shown in Fig. 2. Considerable IL-12 secretion in the low-dose group was detected, which was significantly higher (P<0.05) than in the other groups in both males and females (Fig. 2A). No statistical difference was found in terms of the release of TNF-α in females or males (Fig. 2B), although a downward trend was observed in males.

**Fig 2.**
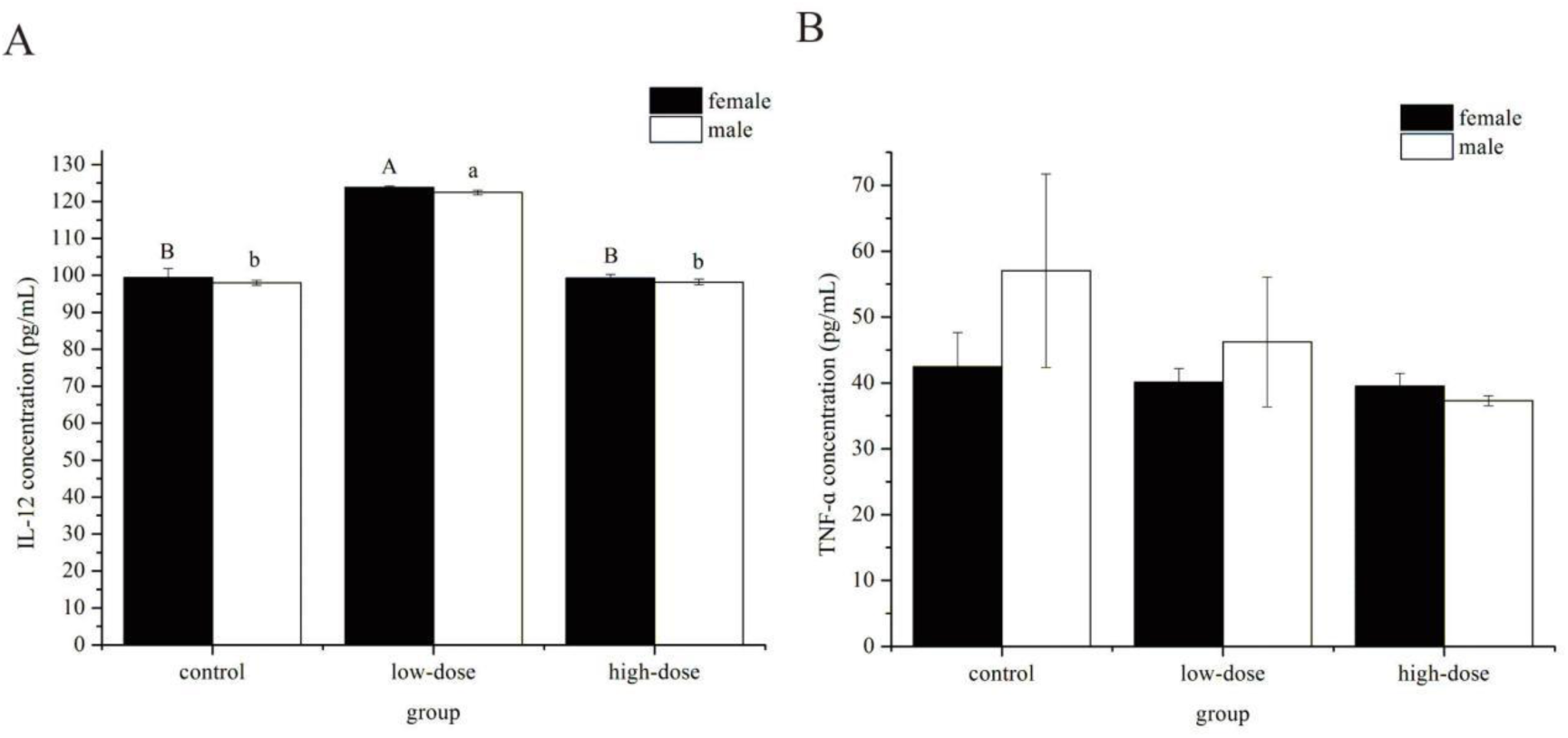
IL-12/TNF-α concentration induced by *L. reuteri* DMSZ 8533. IL-12 concentration induced by *L. reuteri.* Letters in a column mean the indices differed significantly (P<0.05); (B) TNF-α concentration induced by *L. reuteri.* Data are presented as mean ± SE, n =4.

### Immunohistochemistry of ileum section

As Fig. 3A shows, for females, ingestion of low and high doses of *L. reuteri* led to 20.11% and 18.50% increases, respectively, in CD4-positive cells compared with the control group. In contrast, males experienced a 4.42% reduction in such cells after high-dose treatment with *L. reuteri* compared with the control group. Females showed an increase in CD3-positive cells compared with the control group, increasing 23.59% and 13.5% for the low- and high-dose groups, respectively, however, males displayed reductions of 11.25% (low-dose) and 19.5% (high-dose group) compared with the control group (Fig. 3B). An example typical of these findings is shown in Fig. 4B for CD4-posivite cells and in Fig. 4D for CD3-positive cells in the female group treated with low doses of *L. reuteri.* As Fig.5A shows, CD-8 positive cells experienced a increase of 4 % and 3 %, 4.95 % and 1.12 % in females and males, respectively, after ingestion of low and high dose of *L. reuteri*. Furthermore, there was a statistically difference between sexes at high-dose group (P=0.023).

**Fig 3.**
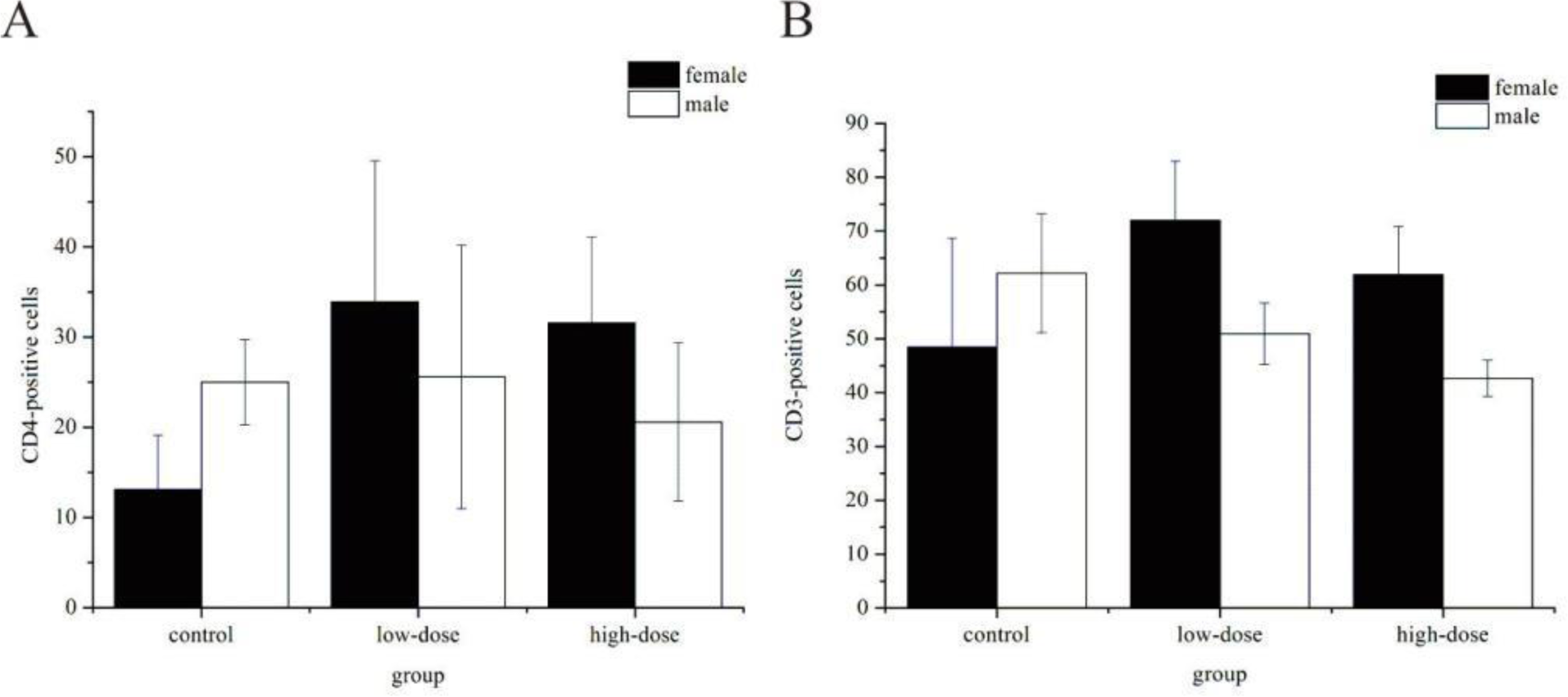
Effect of *L. reuteri* DMSZ 8533 on CD4- and CD3-positive cells in the ileum of BALB/C mice. (A) CD4-positive cells induced by different doses of *L. reuteri*; (B) CD3-positive cells induced by different doses of *L. reuteri.* Data are presented as mean ± SE, n=4.

**Fig 4.**
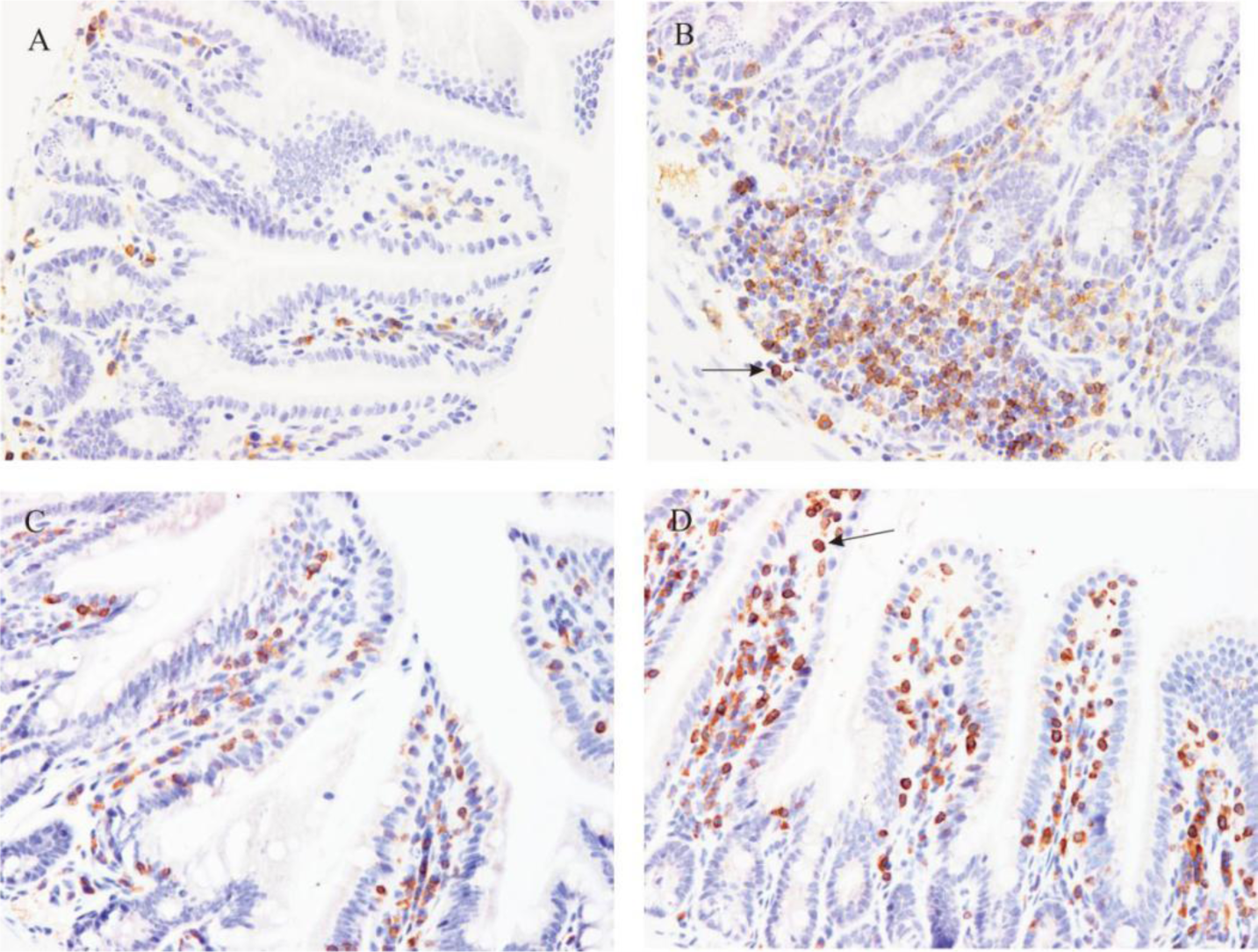
CD4- and CD3-positive cells in female ileal mucosa before and after administration of low doses of *L. reuteri*. (A) and (B), control and low-dose groups for CD4-positive cells; (C) and (D), control and low-dose groups for CD3-positive cells. The arrows in Fig. 4B and Fig. 4D pointed to CD4-positive cells and CD3-positive cells respectively.

**Fig 5.**
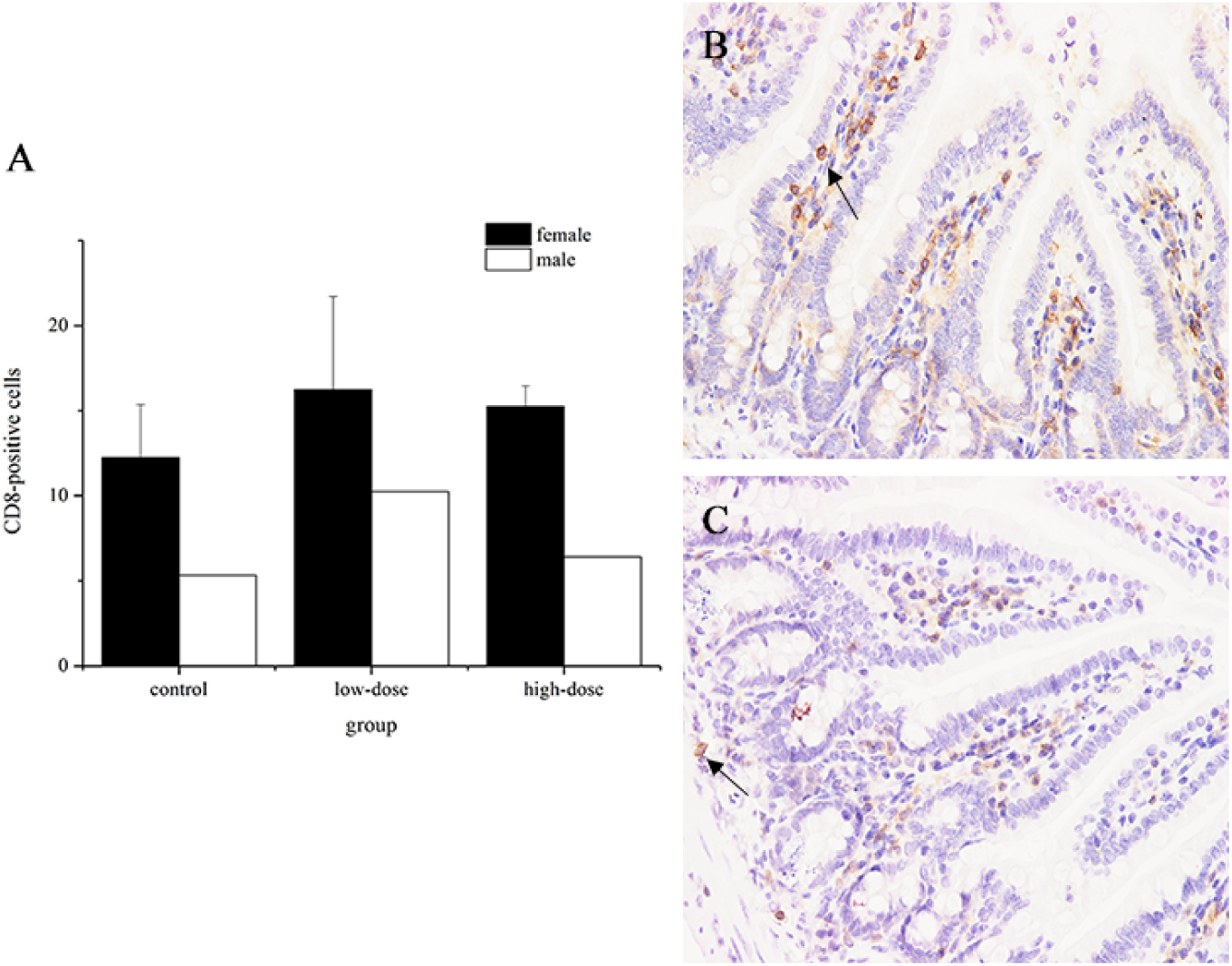
Effect of *L. reuteri* DMSZ 8533 on CD8-positive cells in the ileum of BALB/C mice. (A) CD8-positive cells induced by different doses of *L. reuteri*; (B) CD8-positive cells in female ileal mucosa induced by high doses of *L. reuter*i. (C) CD8-positive cells in male ileal mucosa induced by high doses of *L. reuteri*. Data are presented as mean ± SE, n=4.

### Sex difference on microbial correlation among groups

Using Illumina Miseq platform, a total of 1648603 raw sequences were generated from DNA isolated from fecal samples, and 1551889 valid sequences were filtered out and obtained after chimeras (Table S1). For Alpha diversity analysis (Table S2), high-dose of *L. reuteri* significantly reduced shannon index compared with low-dose group in males, together with an apparent difference between sex (Table S3). While no apparent difference found in females. Based on beta diversity analysis, UniFirac clustering analysis suggested that there were no obvious phylogenetic differences and similar community structure between the control and high-dose groups in females (Fig. 6A). However, the control group results clustered together more closely with males in the low-dose group (Fig. 6B).

**Fig 6.**
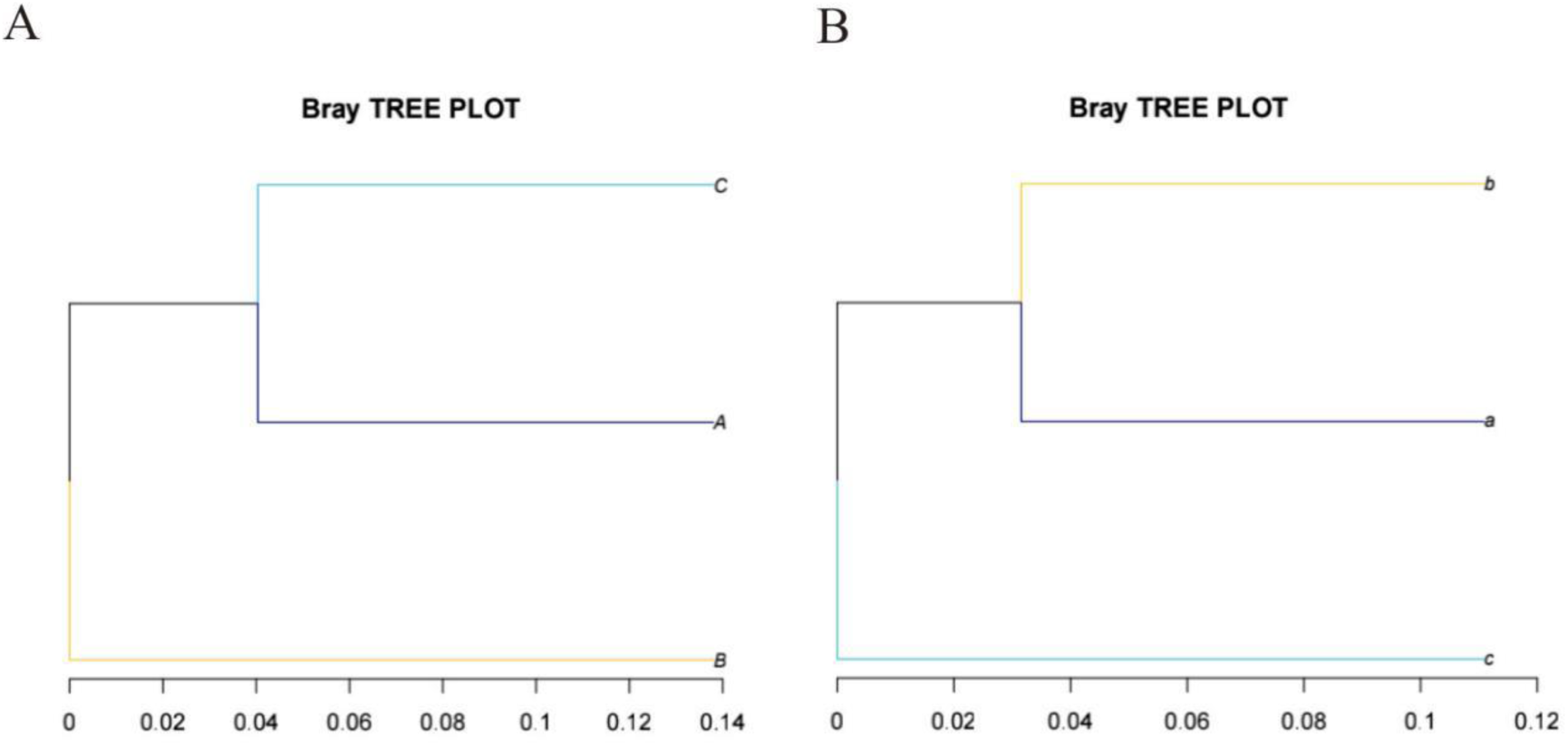
Dose-dependent effect of *L. reuteri* DMSZ 8533 on microflora structure. (A) and (B) Clustering analysis of bacterial community in female and male mice. “a” and “A” refer to control group, “b” and “B” refer to low-dose group, “c” and “C” refer to high-dose group.

### Sex difference on microflora richness at taxonomic level

Taxonomic assignment at the phylum level indicated that gut microbiota shared a similar structure in female and male subjects consisting of four major phyla: *Bacteroidetes*, *Firmicutes*, *Proteobacteria*, and *Actinobacteria*. As Table 2 shows, a constant decline in the abundance of *Actinobacteria* in females was also found in males, but there was a statistically significant sharp decrease in females in the high-dose group. Additionally, *L. reuteri* treatment increased content of phylotypes in the phylum *Bacteroidetes* but decreased *Firmutes,* and opposite observation found in males. At the genus level, relative abundance of *Bacteroides* and *Prevotella* increased in females after *L. reuteri* intervened, while a statistically significant decrease was observed in males. The relative abundance of *Clostridium IV* displayed an reduction in females, while a statistically increase in males. *Lactobacillus* abundance in females was greater than the corresponding groups of males, this appeared to be a concentration-dependent relationship. What’ more, an apparent difference was found between sexes at low-dose group (Table S4). An Ascend trend was found in high-dose group of females in relation to *Lactococcus* abundance while a decrease in males.

**Table 2.**
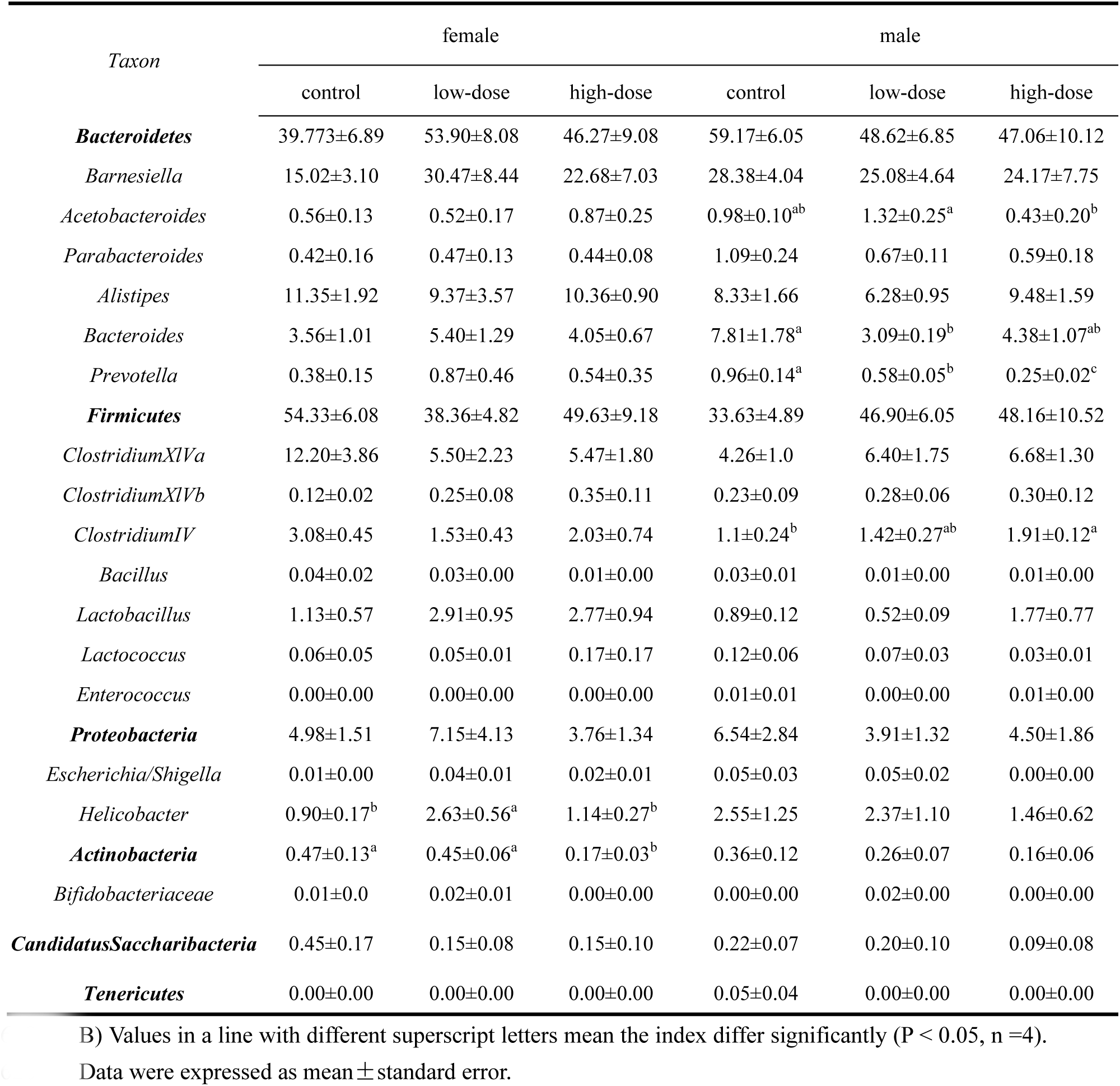
The effect of *L. reuteri* in BALB/c mice at taxonomy level

| Taxon | female |  |  | male |  |  |
| --- | --- | --- | --- | --- | --- | --- |
|  | control | low-dose | high-dose | control | low-dose | high-dose |
| ***Bacteroidetes*** | 39.773±6.89 | 53.90±8.08 | 46.27±9.08 | 59.17±6.05 | 48.62±6.85 | 47.06±10.12 |
| *Barnesiella* | 15.02±3.10 | 30.47±8.44 | 22.68±7.03 | 28.38±4.04 | 25.08±4.64 | 24.17±7.75 |
| *Acetobacteroides* | 0.56±0.13 | 0.52±0.17 | 0.87±0.25 | 0.98±0.10<sup>ab</sup> | 1.32±0.25<sup>a</sup> | 0.43±0.20<sup>b</sup> |
| *Parabacteroides* | 0.42±0.16 | 0.47±0.13 | 0.44±0.08 | 1.09±0.24 | 0.67±0.11 | 0.59±0.18 |
| *Alistipes* | 11.35±1.92 | 9.37±3.57 | 10.36±0.90 | 8.33±1.66 | 6.28±0.95 | 9.48±1.59 |
| *Bacteroides* | 3.56±1.01 | 5.40±1.29 | 4.05±0.67 | 7.81±1.78<sup>a</sup> | 3.09±0.19<sup>b</sup> | 4.38±1.07<sup>ab</sup> |
| *Prevotella* | 0.38±0.15 | 0.87±0.46 | 0.54±0.35 | 0.96±0.14<sup>a</sup> | 0.58±0.05<sup>b</sup> | 0.25±0.02<sup>c</sup> |
| ***Firmicutes*** | 54.33±6.08 | 38.36±4.82 | 49.63±9.18 | 33.63±4.89 | 46.90±6.05 | 48.16±10.52 |
| *ClostridiumXIVa* | 12.20±3.86 | 5.50±2.23 | 5.47±1.80 | 4.26±1.0 | 6.40±1.75 | 6.68±1.30 |
| *ClostridiumXIVb* | 0.12±0.02 | 0.25±0.08 | 0.35±0.11 | 0.23±0.09 | 0.28±0.06 | 0.30±0.12 |
| *ClostridiumIV* | 3.08±0.45 | 1.53±0.43 | 2.03±0.74 | 1.1±0.24<sup>b</sup> | 1.42±0.27<sup>ab</sup> | 1.91±0.12<sup>a</sup> |
| *Bacillus* | 0.04±0.02 | 0.03±0.00 | 0.01±0.00 | 0.03±0.01 | 0.01±0.00 | 0.01±0.00 |
| *Lactobacillus* | 1.13±0.57 | 2.91±0.95 | 2.77±0.94 | 0.89±0.12 | 0.52±0.09 | 1.77±0.77 |
| *Lactococcus* | 0.06±0.05 | 0.05±0.01 | 0.17±0.17 | 0.12±0.06 | 0.07±0.03 | 0.03±0.01 |
| *Enterococcus* | 0.00±0.00 | 0.00±0.00 | 0.00±0.00 | 0.01±0.01 | 0.00±0.00 | 0.01±0.00 |
| ***Proteobacteria*** | 4.98±1.51 | 7.15±4.13 | 3.76±1.34 | 6.54±2.84 | 3.91±1.32 | 4.50±1.86 |
| *Escherichia/Shigella* | 0.01±0.00 | 0.04±0.01 | 0.02±0.01 | 0.05±0.03 | 0.05±0.02 | 0.00±0.00 |
| *Helicobacter* | 0.90±0.17<sup>b</sup> | 2.63±0.56<sup>a</sup> | 1.14±0.27<sup>b</sup> | 2.55±1.25 | 2.37±1.10 | 1.46±0.62 |
| ***Actinobacteria*** | 0.47±0.13<sup>a</sup> | 0.45±0.06<sup>a</sup> | 0.17±0.03<sup>b</sup> | 0.36±0.12 | 0.26±0.07 | 0.16±0.06 |
| *Bifidobacteriaceae* | 0.01±0.0 | 0.02±0.01 | 0.00±0.00 | 0.00±0.00 | 0.02±0.00 | 0.00±0.00 |
| ***CandidatusSaccharibacteria*** | 0.45±0.17 | 0.15±0.08 | 0.15±0.10 | 0.22±0.07 | 0.20±0.10 | 0.09±0.08 |
| ***Tenericutes*** | 0.00±0.00 | 0.00±0.00 | 0.00±0.00 | 0.05±0.04 | 0.00±0.00 | 0.00±0.00 |

B) Values in a line with different superscript letters mean the index differ significantly (P < 0.05, n =4).
Data were expressed as mean ± standard error.

## Discussion

Tolerance to the stressful conditions found in the GIT and adherence to epithelial cells are commonly believed to be key factors in identifying probiotics [31]. The pH in the human stomach is approximately 2.0 to 2.5 [32] and the bile content in the upper small intestine ranges from 0.3% to 0.5% [33,34], which can destroy *Lactobacillus* and the attendant potential probiotic characteristics. Adhesion is another essential criterion to select a potential probiotic strain, since it is related to the residence time of *Lactobacillus* and, possibly, temporal colonization [35]. In our study, *L. reuteri* displayed promising survival rate and adhesion ability values, which is consistent with another study [36]. Based on these results, we conducted *in vivo* experiments to evaluate the effect of this possibly probiotic *L. reuteri* strain on immune response and microflora diversity.

Recently, mocusal immunity mediated by *L. reuteri* has sparked increased interest among researchers [37,38]. T lymphocytes initiate their helper or cytotoxic roles when they are activated by antigens or pathogens. In our study, low and high doses of *L. reuteri* induced a considerable increase in the number of T lymphocytes in females, which agreed with previous observations [37]. In contrast with changes in females, a slight decline was obtained in CD4- and CD3-positive cells in males. We hypothesized that the difference was due to sex hormones, such as estrogen or 17ß-estradio (E2), which are widely accepted as enhancers of CD4^+^ T cell expansion [39,40]. In mice, there are at least three CD4^+^ subsets: Th1, Th2 and Th0. Th1 mainly mediate cellular immunity associated with the secretion of the cytokines IL-12, IFN-γ and TNF-α. IL-12, produced by cells as part of the innate defense system in response to bacteria, is primarily IFN-γ [41]. Low doses of *L. reuteri* were the most significant inducer of IL-12 in males and females, confirming that *L. reuteri* possess potent immune-enhancing properties. Our findings were consistent with other findings [42]. However, high doses of *L. reuteri* had no effect on IL-12 secretion compared with the control group, which also agreed with previous research [43] that *Lactobacillus* were active only at low-bacterial concentrations. This finding might be ascribed to low and high dose microbe-associated molecular patterns (MAMP) mediating different receptor conformations, or they may differentially distribute to subcellular locations, subsequently activating different downstream pathways [44]. We further investigated the level of TNF-α, a major pro-inflammatory cytokine that can mediate the inflammatory response at a systemic level [45]. Notably, concentrations of TNF-α in males was much higher than that found in females. The higher content of *Helicobacter* and *Escherichia* in males (Table 2) might support this finding, which can both up-regulate TNF-α [46]. Furthermore, the TNF-α concentration exhibited a downward trend as *L. reuteri* concentration increased in males, possibly due to suppression by *L. reuteri* through the conversion of histamine via PKA and ERK signaling [47].

The intestinal microflora constitute a commensal anaerobic and facultative microorganism, which seeks to inhibit pathogens, participates in anabolic pathways and maintains immune hemostasis [48]. Many genera belonging to *Bacteroidetes*, such as *Barnesiella* and *Bacteroides*, could modulate the immune response by enhancing populations of marginal zone B cells, natural killer T cells [49] or capsular polysaccharide biosynthesis [50]. In the present study, *L. reuteri* treatment increased the abundance of phylum *Bacteroidetes* (primarily the genus of *Barnesiella, Bacteroides*) in females but not in males. Another dominant phylum, *Firmicutes,* mainly consisted of the *Clostridia* class and *Bacilli class*, which are capable of producing lactic acid and carbon dioxide [51]. In our study, we observed a reserve shift in the content of *Firmicutes* between sexes after *L. reuteri* treatment, which occurred largely relevant with the opposite variation of the phylum *Bacteroidetes* content. Additionally, the abundance of genus *Bacillus* and *Lactobacillus*, which both belonging to class *Bacilli*, also statistically different in the sexes, especially in the low-dose group (Table S4). However, *Lactobacillus* are generally considered immuno-mediators in the human GIT and can mediate mucosal immunity [37]. What’ more, the antagonistic effect of *Lactobacillus* was also illustrated in our present study, with the abundance of *Lactobacillus* increased and the amount of harmful bacteria, such as *Enterococcus, Helicobacter* and *Escherichia* suppressed, consistent with Xie et al’s findings [52]. The antagonistic actions of *Lactobacilli* might be the result of a pH reduction [53], bacteriocin secretion [54] and/or competition for adhesion sites with harmful bacteria. *Clostridial spp*, such as *Clostridium IV*, which populates the ileum and cecum in mice, can produce short-chain fatty acids and induce Treg cells [55]. Positive correlations between the abundance of *Clostridium IV* and the concentration of *L. reuteri* were found in males while a reduction phenomenon in females. However, the abundance of *Clostridium IV* in females was greater than that in males in the corresponding dose groups.

## Conclusion

As the most resistant probiotic strain found in our *in vitro* experiments, *L. reuteri* DMSZ 8533’s effect on the immune parameters of, and microbial composition in, BALB/c mice were affected by sex differences. Our results demonstrated that *L. reuteri* exerted a positive effect on the immune response in females, with higher levels of T lymphocytes and IL-12 observed at low doses of *L. reuteri*. However, males did not exhibit the same response, except in relation to IL-12 and CD-8 T cells. In addition, *L. reuteri* treatment exihibited a reserve shift between sexes on the content of phylotypes in the phylum *Bacteroidetes* (genus of *Bacteroides* and *Prevotella) and Firmicutes* (genus of *ClostridiumIV*).

## Materials and Methods

*Lactobacillus reuteri* DMSZ 8533 was purchased from the China Center of Industrial Culture Collection (Beijing, China). *Lactobacillus plantarum* 14917 was preserved in our laboratory, while *Lactobacillus acidophilus* ATCC 4356 was obtained from the China General Microbiological Culture Collection Center (Beijing, China). Bacteria were incubated anaerobically in MRS broth at 37 °C for 24 h before use. Bacteria were stored at −80 in 30% (v/v) glycerol.

Caco-2 cells obtained from the Biological Technology Co. (California, USA), which isolated from human adenocarcinoma of colon, were grown in DMEM (Sigma, Darmstadt, Germany) containing 20% (v/v) fetal bovine serum (FBS) and 1% (v/v) antibiotics (100 U/mL penicillin, 100 mg/mL streptomycin) at 37 °C in a 5% CO2 atmosphere. For the adhesion assay, 2 mL of cells were seeded in a 6-well tissue culture plate without antibiotics and cultivated until 90% confluence was achieved.

### Gastrointestinal tolerance analysis

Simulated gastric digestion was performed as described previously [56] with slight modifications. Overnight cultures (4 mL) were harvested by centrifugation at 6000×g for 10 min and suspended in 4 mL artificial gastric juice adjusted to pH 2.0 with 1 mol/L hydrochloric acid. The bacterial suspension was incubated for 3 h at 37 °C. The aliquots were taken for the enumeration of viable cells at 0 and 180 min. After treatment with artificial gastric juice, bacteria were collected by centrifugation (6000×g, 10 min) and suspended in simulated intestinal fluids adjusted to pH 8.0 with 1 mol/L NaOH for another 4 h. The colony forming unit (CFU) enumeration was determined at 0 and 4 h.

### Auto-aggregation analysis

Auto-aggregation assay was assessed according to the method described by Xu et al [57]. Briefly, 5 mL bacteria cells in the stationary phase were centrifuged, washed twice with 0.01 M phosphate buffered solution (PBS) and re-suspended in PBS to OD_600_=0.5±0.02 (A_0_). Then 3 mL bacterial suspension was vortexed for 15 s and incubated at room temperature for 3 h. Then 0.4 mL of the upper suspension was removed to determine the absorbance (A_1_) at 600 nm. The auto-aggregation percentage was expressed as:

Auto-aggregation rate (%) = (1-A_1_/A_0_) × 100

### Cell-surface hydrophobicity analysis

Microbial adhesion to solvents (MATS) was used to evaluate the surface hydrophobicity property of the Lactobacillus with slight modifications [58,59]. Bacteria in the stationary phase were harvested, washed twice with PBS, and re-suspended in 0.1 M KNO_3_ to 10^8^ CFU/mL (A_0_). Then 1.2 mL of bacteria suspension was mixed with 0.2 mL xylene and vortexed for 2 min after achieving a stationary condition for 10 min. To ensure the complete separation of the mixture, the aqueous phase was removed after 30 min of incubation at room temperature and its absorbance (A_1_) was measured at OD_600_. The percentage of microbial adhesion to solvents was calculated as:

Cell surface hydrophobicity (%) = (1-A_1_/A_0_) × 100.

### Adhesion analysis

Bacteria in the stationary phase were centrifuged (6000×g, 10 min), washed twice and re-suspended in PBS adjusted to OD_600_=0.7±0.05. Then they were mixed with 10 μM 6-carboxyfluorescein diacetate (CFDA; Sigma, Darmstadt, Germany) and incubated at 37 °C for 30 min. Bacteria were washed three times to remove any unmarked CFDA. Caco-2-coated wells were washed twice with PBS, and 2 mL of CFDA-labelled bacteria were added. The mixture was incubated in a 5% CO2 atmosphere at 37 °C for 2 h. After incubation, the wells were washed three times with 2 mL PBS to remove the un-adhered bacteria, and then 1.4 mL 0.25% EDTA-trypsin was added into the wells for 10 min. Then 0.6 mL DMEM was added to terminate digestion. Finally, fluorescence intensity (excitation wavelength 485 nm; emission wavelength 538 nm) was measured using a Tecan Infinite M200 Pro (Tecan Group, Switzerland) and the adhesion rate was expressed as the percentage of fluorescence recovered after binding to Caco-2 cells relative to the fluorescence of the bacterial suspension added to the wells.

### Animals and treatment

BALB/c mice (6 weeks old) purchased from the Zhejiang Academy of Medical Sciences (Hangzhou, China) were housed in an automatic light/dark cycle (light periods of 12 h) habitat, and provided water and rodent chow during the whole experiment under specific pathogen-free (SPF) conditions (Laboratory Animal Center of Ningbo University). Mice were randomly divided into three groups (n=12/group, 6 each sex): control and low- and high-dose groups. For low- and high-dose groups, mice received 0.4 mL of skimmed milk containing 10^8^ CFU/mL or 10^10^ CFU/mL *L. reuteri* DMSZ 8533, respectively, by oral gavage every day for four weeks. Mice in the control group were administrated 0.4 mL of skimmed milk containing no *L. reuteri*. All animal care and experimental procedures were approved by the Committee on Animal Care and Use, and the Committee on the Ethics of Animal Experiments of Ningbo University. Before surgery, mice were weighed and injected with 5 mg/kg Carprophen (Rimadyl) as an analgesic. Then mice were anesthetized (isoflurane 2–3% mixed with 30% oxygen (O_2_) and 70% nitrous oxide (N_2_O) before eyeball extirpating. Then the spleen, thymus and liver were weighed.

### Enzyme-linked immunosorbent assay

Blood drawn from the eyes was collected, kept at room temperature for 1.5 h and centrifuged (2500×g, 30 min). The obtained serum was frozen (−80 °C) for IL-12 and TNF-α determination using a mouse IL-12/TNF-α Elisa Kit (Lianke-Biotech Co., Hangzhou, China).

### Immunohistochemistry

For immunohistochemistry, the method was performed according to previous research [60]. Briefly, ileum sections were fixed in 4% paraformaldehyde (Sigma-Aldrich, USA) and embedded in paraffin. Sections were baked at 60 °C overnight, de-paraffinized in xylene and hydrated in graded ethanol. Antigen retrieval was performed in citrate antigen retrieval solution to maintain pressure for 4 min and cooled to room temperature. Drops of 3% H_2_O_2_ were added to quench any endogenous peroxide activity. Tissue sections were blocked with blocking buffer (Sangon-Biotech, Shanghai, China) for 45 min, rinsed and incubated with properly diluted primary antibody (anti-CD4 or anti-CD8 antibody, Abcam; anti-CD3 antibody, Cell Signaling Technology) overnight. Goat anti-rabbit secondary antibody was then used to detect anti-CD4, anti-CD3 or anti-CD8. Streptavidin-HRP and Diaminobenzidine (DAB) were used to visualize regions of tissue with anti-CD4, anti-CD3 or anti-CD8. Hematoxylin was used to counter-stain cells. The CD-8, CD4- and CD3-positive cells were counted in three fields using an Olympus BX53 (Pennsylvania, USA) microscope at 400 ×magnification.

### 16S rRNA amplification and MiSeq sequencing

The V3-V4 variable region of the 16S rRNA gene was amplified from 35 fecal DNA extracts using the 16S meta-genomic sequencing library protocol (Illumina). Two rounds of PCR amplification were completed on the fecal DNA. The initial step was performed with the PCR primers targeting the V3-V4 region of the 16sRNA gene: 341F and 805R (Forward primer: CCCTACACGACGCTCTTCCGATCTGCCTACGGGNGGCWGCAG; Reverse primer: GACTGGAGTTCCTTGGCACCCGAGAATTCCA). All PCR reactions were performed with Ex Taq ploymerase (Takara, Shanghai, China) with approximately 10-20 ng of genomic DNA as follows: heated lid at 94° C for 3 min; followed by 5 cycles of amplification at 94 °C (30 s), 45 °C (20 s) and 60 °C (30 s); 20 cycles of amplification at 94 °C (20 s), 55 °C (20 s) and 72 °C (30 s); and a final extension step of 5 min at 72 °C. Successful amplicons were purified with an Easypure quick gel extraction kit (Transgen Biotech; Beijing, China). A second round of amplification was performed with 20 ng of purified DNA and primers containing the Illumina adapters and indexes. PCR cycling conditions were as follows: 95 °C for 3 min; 5 cycles at 94 °C for 20 s, 72 °C 30 s, 72 °C 30 s; and a final extension step at 72 °C for 5 min. All purified DNA amplicons were pooled in equimolar concentrations and the final concentration was adjusted to 20 pmol for subsequent metagenomic sequencing.

### Bioinformatics and statistical analysis

Data were analyzed using a one-way analysis of variance (ANOVA) and independent-samples test using 16.0 SPSS software (SPSS Inc., USA) with P values of 0.05 considered significant.

For gut microbiota analysis, raw sequences were trimmed to remove sequence of primer joint, and high-quality pair-end reads were merged on tags through overlaps using Pear (v 0.9.6) software. The valid data were obtained through distinguished by unique barcode sequences, filtered by quality control and remained after chimeras. The tag were clustered to operational taxonomic units (OTUs) using Ultra-fast sequence analysis (Usearch version 5.2.236) with 97 % identity level. Relative abundance of bacterial taxa were determined for each community by comparing the number of reads assigned to a specific taxa to total number of reads. Alpha diversity used to analyze the species diversity and richness was calculated by Mothur (version 1.30.1) software. Bray-Crutis trees based on genus level was carried out by Unweighted Pair Group Method with Arithmetic mean (UPGMA) and was done with software programme R(version 3.2). UniFrac, including weighted UniFrac and unweighted UniFrac, used the systematic evolution information to compare the composition of the microbial community between samples.

## Acknowledgement

This work was supported by the Natural Science Funding of China (31601487, 31671869 and 31471598), the Science and Technology Bureau of Ningbo (2016C10022), and the K. C. Wong Magna Fund in Ningbo University.

